# Nutrient enrichment alters gene expression in ‘*Ca.’* Aquarickettsia rohweri, promoting parasite expansion and horizontal transmission

**DOI:** 10.1101/2023.06.08.544262

**Authors:** Lauren Speare, J Grace Klinges, William C Duke, Erinn M Muller, Rebecca L Vega Thurber

## Abstract

Ocean warming, disease, and pollution contributed to global declines in coral abundances and diversity. In the Caribbean, corals previously dominated reefs, providing an architectural framework for diverse ecological habitats, but have significantly declined due to infectious microbial disease. Key species like coral *Acropora cervicornis*, are now considered critically endangered, prompting researchers to focus on scientific endeavors to identify factors that influence coral disease resistance and resilience. We previously showed that disease susceptibility, growth rates, and bleaching risk were all associated with the abundance of a single bacterial parasite, ‘*Ca.’* Aquarickettsia rohweri which proliferates *in vivo* under nutrient enrichment. Yet how nutrients influence parasite physiology and life history strategies within its host are unknown. We performed microscopy and transcriptomic analyses of ‘*Ca.’* A. rohweri populations during a 6-week nutrient exposure experiment. Microscopy showed that this parasite was abundant in coral tissue and densely packed in mucocytes prior to nutrient enrichment. ‘*Ca.’* A. rohweri energy scavenging genes and those potentially involved in this habitat transition are significantly upregulated during enrichment. Specifically, transcripts involved in signaling, virulence, two-component systems, and nutrient import genes are elevated under higher nutrients. These data support the predicted role of ‘*Ca.’* A. rohweri as a highly active nutrient-responsive *A. cervicornis* parasite, and provide a glimpse at the mechanism of induced disease susceptibility while implicating nutrient exposure in its horizontal transmission.

**Significance:** The coral disease crisis has contributed to global declines in coral abundance and diversity and is exacerbated by environmental stressors like eutrophication. Thus, identifying factors that influence coral disease resistance and resilience is a top priority. The Rickettsiales-like bacterium, *‘Candidatus’* Aquarickettsia rohweri is ubiquitous coral symbiont that is strongly linked to coral disease susceptibility in staghorn coral, and is undergoing positive selection across the Caribbean. Although *‘Ca.’* A. rohweri is a putative parasite, little is known about the activity of this bacterium in coral tissue. This work supports the role of *‘Ca.’* A. rohweri as a highly active, nutrient-responsive parasite and proposes a mechanism for how *‘Ca.’* A. rohweri contributes to coral disease susceptibility, parasite expansion, and horizontal transmission.

## Introduction

Environmental stressors such as anthropogenic-induced ocean warming, disease, and pollution have contributed to a world-wide decline in coral diversity and coverage (1, 2). Corals maintain important associations with a myriad of microbial symbionts; however, these intricate relationships can be disrupted by environmental disturbances, resulting in dysbiosis and coral disease (3–9). For example, the relationship between corals and their endosymbiotic dinoflagellates, which provide corals with sugars and essential amino acids, is dependent on oligotrophic conditions with low nitrogen availability to promote phosphorus cycling (10, 11). Local eutrophication has disrupted this delicate balance, resulting in increased prevalence and severity of coral bleaching and disease (7, 12, 13). In the Caribbean, Acroporid corals that previously dominated reefs and provided the architectural framework for diverse ecological habitats have shown significant declines due to infectious disease (14–17). Species like the staghorn coral *Acropora cervicornis*, are some of the only fast-growing taxa with branching morphologies in the region and are now considered critically endangered. This has prompted researchers to focus restoration efforts on understanding factors that promote disease susceptibility and resistance.

Recent evidence suggests that host genotype and microbiome composition significantly impact *A. cervicornis* disease susceptibility (18–21). Disease resistant hosts may better tolerate potential pathogens, prevent opportunists from acting antagonistically, or house beneficial symbionts that increase host disease resistance (21). In contrast, susceptible genotypes may more easily succumb to microbial antagonism or harbor parasites that exacerbate environmental stressors. For example, *A. cervicornis* disease susceptibility was recently linked to the presence of an intracellular bacterial parasite, ‘*Candidatus’* Aquarickettsia rohweri (18, 22, 23). It was found that ‘*Ca.’* A. rohweri abundance also varies significantly with *A. cervicornis* genotype, where microbiomes of disease susceptible genotypes are dominated by *Ca. A. rohweri* (89.7%), while ‘*Ca.’* A. rohweri makes up a minor constituent of disease resistant microbiomes (2.5%) (18, 19, 24). Further, ‘*Ca.’* A. rohweri abundance was experimentally linked to reduced coral growth rates (9) and increased infection by opportunists upon bleaching (18).

It was also recently shown that ‘*Ca.’* A. rohweri are undergoing positive selection across the Caribbean and most strongly in Florida, suggesting they are highly responsive to their environment (25). Speciation and virulence genes, including type IV secretion system (T4SS) genes, are undergoing the greatest degree of positive selection, which is concerning given that ‘*Ca.’* A. rohweri are also transmitted horizontally between hosts (25). Like other Rickettsiales bacteria (26), ‘*Ca.’* A. rohweri appears to parasitize its host for nutrients, energy, and amino acids (22). ‘*Ca.’* A. rohweri inhabit coral mucocytes, cells in the coral epidermis that produce mucus to protect against sedimentation and infection; within these specialized cells, ‘*Ca.’* A. rohweri are localized near, yet not within, endosymbiotic coral dinoflagellate cells (22). This parasite does not encode genes to synthesize most amino acids (22), suggesting it relies heavily on parasitism of the coral host and/or endosymbiotic dinoflagellates (18) for resources.

Given the pervasiveness of ‘*Ca.’* A. rohweri in Caribbean Acroporids and demonstrated negative effects of local nutrient pollution on coral health, significant efforts have gone towards understanding the role of nutrient enrichment on ‘*Ca.’* A. rohweri abundance (9, 27). Recently, we performed a manipulative tank experiment to disentangle the individual and combined impacts of eutrophication on ‘*Ca.’* A. rohweri abundance within *A. cervicornis* microbiomes (24, 27). We exposed a disease susceptible *A. cervicornis* genotype (ML-50) and a disease resistant *A. cervicornis* genotype (ML-7), to elevated levels of nitrate, ammonium, phosphate, or a combination of the three for six weeks and evaluated microbiome composition and host fitness. Analysis of community dynamics (i.e., 16S amplicon libraries) and absolute abundance (quantitative polymerase chain reaction (qPCR) of a ‘*Ca.’* A. rohweri specific marker gene) revealed that ‘*Ca.’* A. rohweri responded positively to nutrient enrichment in both host genotypes, however it remained at low relative abundances in the disease-resistant genotype, ML-7 (<0.5% of the total bacterial community) (24). In the disease-susceptible genotype, M-50, *‘Ca.’* A. rohweri dominated the bacterial microbiome in all treatments and increased in both relative and absolute abundance, while overall microbiome diversity declined in response to nutrient enrichment (27). Genotype ML-50 Corals showed an increase in visual dinoflagellate symbiont density yet a decrease in coral growth in response to elevated nutrients, suggesting that nutrient enrichment promotes coral microbiome dysbiosis and reduced coral fitness (27). Given the increase in ‘*Ca.’* A. rohweri abundance and dominance within the microbiomes of *A. cervicornis* genotype ML-50, we hypothesized that inorganic nutrient enrichment also increases parasitic activity through transcription of energy scavenging genes, thus weakening and eventually killing the host during additional disturbances such as temperature stress.

In this work, we tested the hypothesis that nutrient enrichment promotes parasitic gene expression by performing meta-transcriptomic and transmission electron microscope analyses of ‘*Ca.’* A. rohweri populations within holobiont tissue. Aquarium conditions in our unamended nutrient treatment had moderately elevated nutrient concentrations compared to the coral collection site, yet lower concentrations than the combined nitrate, ammonium, phosphate enrichment treatment (Fig S1) (27). Thus, by comparing samples collected at the beginning of the experiment, to samples maintained in our “unamended/baseline” and “nutrient enriched” treatments for six weeks, we could examine ‘*Ca.’* A. rohweri gene expression across a nutrient gradient. Here, we describe the localization and transcriptional activity of ‘*Ca.’* A. rohweri *in vivo* in a disease-susceptible *A. cervicornis* genotype (ML-50), show evidence supporting the predicted role of ‘*Ca.’* A. rohweri as a nutrient-responsive *A. cervicornis* parasite, and provide a glimpse as to how this widespread microbe contributes to increased coral disease susceptibility and mortality.

## Results

### ‘*Ca.’* A. rohweri is prevalent in *A. cervicornis* genotype ML-50 mucocytes and tissue

To test our hypothesis that nutrient enrichment affects expression of key genes and ‘*Ca.’* A. rohweri symbiosis status *in vivo*, we analyzed tissue samples and metatranscriptomes of *A. cervicornis* genotype ML-50 collected from a previous nutrient enrichment experiment (27). Briefly, *A. cervicornis* fragments were collected from Looe Key, allowed to acclimate to aquarium conditions for seven days, and placed into one of two experimental treatments for six weeks: 1) baseline aquarium conditions that had 4x offshore reef nitrate concentrations, or 2) ‘nutrient amended/enriched’ that had 12-16x offshore reef concentrations of nitrate (Na_2_NO_3_), ammonium (NH_4_Cl) and phosphate (Na_3_PO_4_) (Fig S1). We first examined the localization and distribution of ‘*Ca.’* A. rohweri cells within coral tissue prior to nutrient enrichment via scanning electron microscopy (SEM) (Fig 1). Similarly to previous observations (22), Rickettsiales-like organisms (RLOs) were prevalent within coral tissues, both outside of coral mucocytes and also densely packed within mucocytes (Fig 1). These Rickettsiales filled mucocytes (RFMs) were abundant within both gastrodermal cells and within the epidermis. Rickettsiales cells found inside RFMs were ∼1-2.5 um in length and 0.5 um in width. Given that 99.9% of the bacterial community of these corals is ‘*Ca.’* A. rohweri, we can assume a majority of these packaged cells are ‘*Ca.’* A. rohweri. The presence of these densely packaged RFMs suggests that one mechanism of parasite horizontal transmission is through infected mucocyte release into the surrounding water column.

### Experiment duration and nutrient enrichment shifts ‘*Ca.’* A. rohweri gene expression

To examine the impact of nutrient enrichment on ‘*Ca.’* A. rohweri gene expression *in vivo*, we analyzed the metatranscriptomes of *A. cervicornis* genotype ML-50 and ML-7 tissues. RNA samples were collected at the beginning of the experiment (time zero) and after six weeks. As corals were held in raceways (tanks) on shore which had conditions distinct from their natural habitat - nutrients were elevated in the aquaria system relative to the nursery from which they were collected - during one week of acclimation, we have chosen to define this cohort of time zero samples as ‘Acute exposure to Baseline aquaria nutrient conditions’ (AB) to most accurately represent their experimental nutrient history. Samples that were held in the aquaria for the six week experiment (seven total weeks including acclimation) are defined as either ‘Chronic exposure to Baseline aquaria nutrient conditions’ (CB) or ‘Chronic exposure to Enriched nutrients’ (CE), 12-16x offshore reef concentrations.

**Figure 1.**
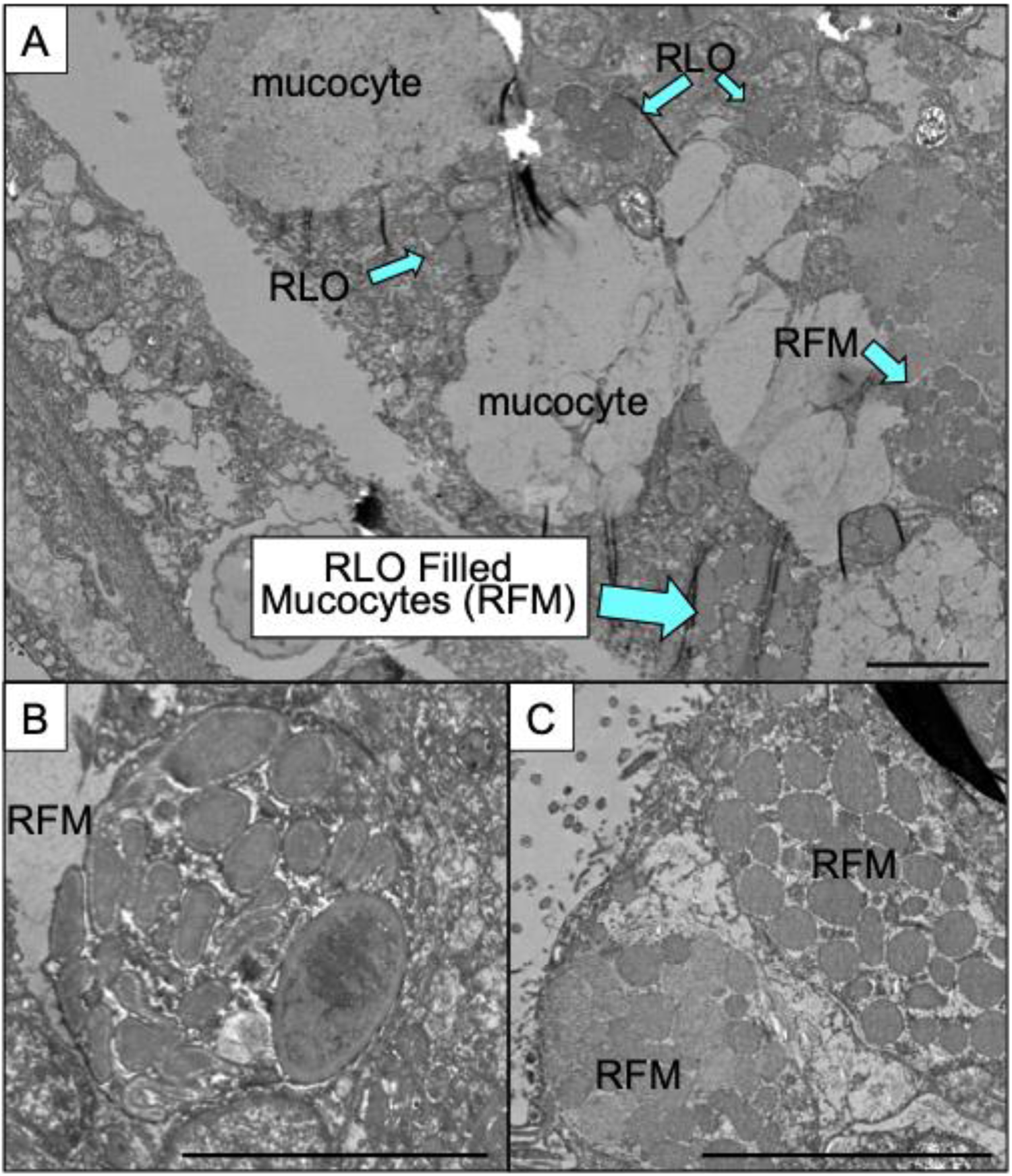
Representative images of *Acropora cervicornis* tissues during experiment. (A) Examples of apparently normal mucocytes and mucocytes filled with Rickettsiales Like Organisms (RLOs) as well as clusters of RLOs outside of mucocytes. (B) Individual RLO Filled Mucocyte (RFM) with multiple bacterial cells. (C) Two side-by-side mucocytes filled with bacterial cells at the edge of the epithelium ready to release infected mucocyte into the environment. Scale bars in all images indicate 5 um.

Transcripts that mapped to the ‘*Ca.’* A. rohweri genome made up approximately 0.1% of the entire ML-50 metatranscriptome for each sample, or roughly 30,000 reads out of ∼35 million reads (Fig S2A) in each sample (Fig S2B). This percentage is comparable to a recent study that showed bacterial transcripts made up ∼0.09% of annotated transcripts for the entire coral metatranscriptomes (28). Transcripts were detected for 65-71% of the coding region of the ‘*Ca.’* A. rohweri genome and were not detected at significantly different levels between experimental treatments (Fig S2C). Although RNA-Seq observations from bacteria maintained in pure cultures indicate that most, if not all, genes are expressed in the bacterial genome given deep enough sequencing (29), achieving such transcriptome coverage for an organism in a complex consortium like the coral holobiont remains challenging. Thus, genes with naturally low levels of transcription may not be included in this analysis. Conversely, we only detected *‘Ca.’* A. rohweri transcripts for less than 0.003% of the ML-7 metatranscriptome, or roughly 1,000 reads out of ∼37 million reads. Transcripts were detected for less than 0.14% of the coding region of the *‘Ca.’* A. rohweri genome (2 genes) in all samples. Because of this minimal amount of *‘Ca.’* A. rohweri transcripts, likely due to the low abundance of this bacterium within *A. cervicornis* genotype ML-7 microbiomes (<0.5%), we were unable to proceed analyzing *‘Ca.’* A. rohweri transcription in this coral genotype.

To begin understanding the impact of nutrient enrichment on ‘*Ca.’* A. rohweri activity in disease-susceptible *A. cervicornis* tissue, we first performed a principal coordinates analysis on our ‘*Ca.’* A. rohweri transcriptomes. PCoA demonstrated that samples clustered according to their treatment, where Acute Baseline (AB), Chronic Baseline (CB), and Chronic Enriched (CE) samples clustered separately from one another (Fig 2A). Experimental treatment explained 28.8% percent of the variability in the dataset. There were no significant differences in dispersion (P=0.292) or distance between centroids (P=0.113) for these datasets (Permutation test for homogeneity of multivariate distances), indicating there was no statistical difference in transcriptome variability between treatments.

**Figure 2.**
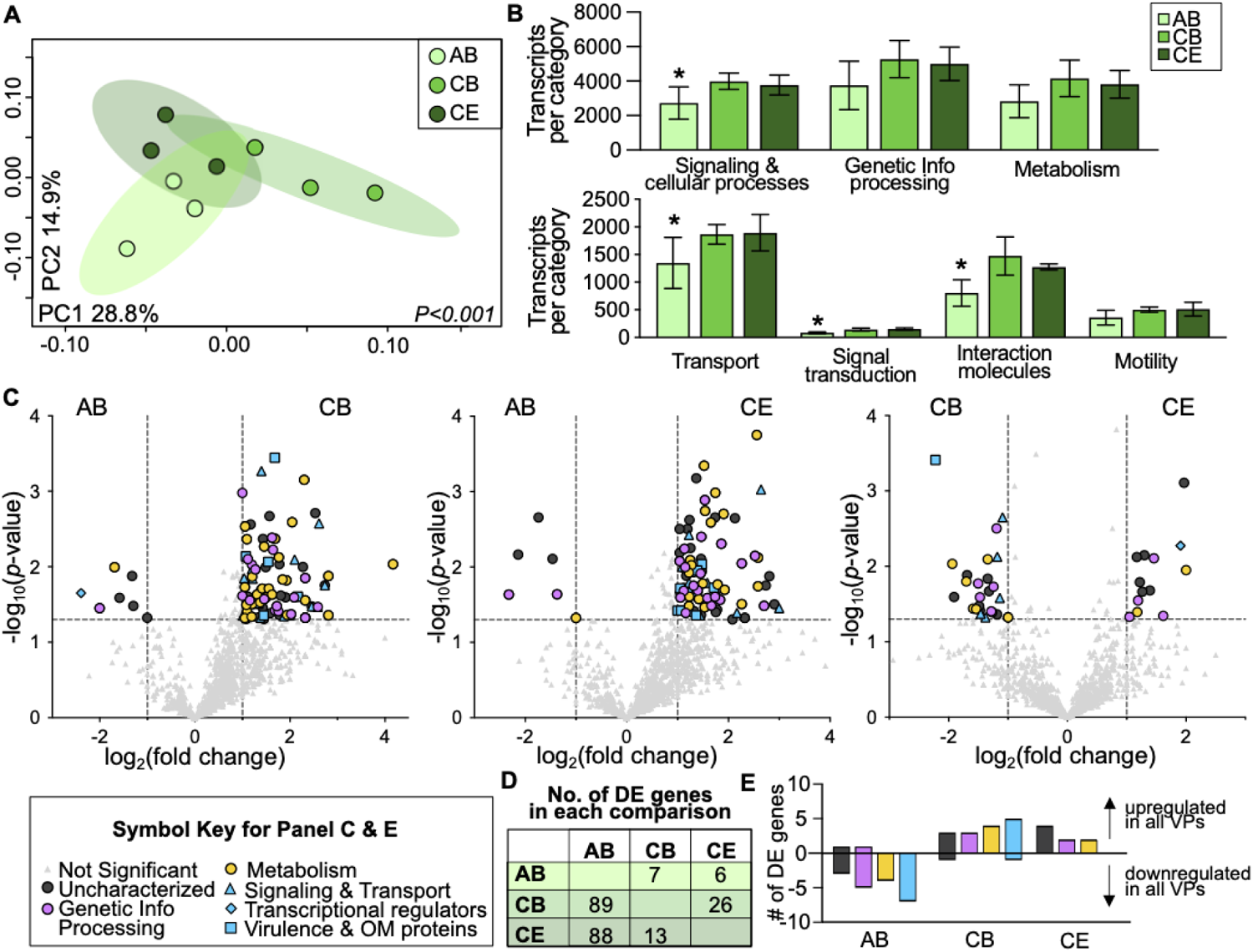
‘*Ca.’* A. rohweri transcriptome is shaped by nutrient enrichment. (A) Principal coordinates analysis (PCoA) based on Bray-Curtis dissimilarities of transcriptomes by nutrient treatment, indicated by symbol color: Acute Baseline (AB, light green), Chronic Baseline (CB, medium green), Chronic Enriched nutrients (CE, dark green). Percentages on each axis indicate the amount of variation explained by each axis; p-value indicates significant results of PERMANOVA tests. (B) Bar charts displaying the number of transcripts for each functional category according to KO designation: broad functional category (top), and specific categories within the signaling & cellular processes category (bottom). Asterisks indicate significantly different numbers of transcripts per category between nutrient treatments (Two-way Analysis of Variance (ANOVA) with Tukey’s multiple comparison post-test, *P*<0.05). (C) Volcano plots showing pairwise comparative analysis of transcript abundance between each treatment. Treatments being compared are shown at the top of each plot. Light gray triangles were not significantly differentially expressed (DE), and all other symbols indicate genes that were significantly DE: a magnitude fold change >|1| log_2_ (vertical dashed lines on x-axis) and p-value <0.05 corrected for multiple comparisons with the Benjamini-Hochberg procedure (horizontal dashed line on y-axis). Symbol color and shape indicate significantly DE genes with KO designation, shown in key. OM indicates ‘outer membrane’. (D) Number (No.) of significantly DE genes in each volcano plot. (E) Number of DE genes that were consistently upregulated (positive values) or downregulated (negative values) in a given nutrient treatment in both volcano plot comparisons (i.e. a gene is significantly upregulated in AB nutrient samples in the AB v CB volcano plot, and the AB v CE volcano plot).

We next sought to determine which functional gene categories were driving the differences between nutrient enrichment levels. Approximately half of the transcribed genes did not have a KO designation and are therefore referred to as “uncharacterized.” The majority of characterized transcripts were ‘genetic information processing’ transcripts, followed by ‘metabolism and environmental information and cellular processes’ which we grouped under the ‘Signaling and Interactions’ category (Fig 2B). There were no significant differences between the percentage of transcripts in each general category across samples within a treatment or across treatments. Therefore, we sought to identify drivers of transcriptome differences at the individual transcript level.

### Signaling genes are disproportionately differentially expressed between acute and chronic treatments

To better understand the transcripts driving the differences between treatments, we generated volcano plots to identify differentially expressed transcripts between acute vs chronic treatments and baseline vs nutrient treatments. Between 8.2 (96 genes) and 8.0% (94 genes) of transcribed genes were differentially expressed (DE) between Acute Baseline and Chronic Baseline or Chronic Enriched nutrient samples, respectively, while only 3.3% (39 genes) of transcribed genes were differentially expressed between Chronic Baseline and Chronic Enriched nutrient samples (Fig 2C). The majority of differentially expressed genes had higher levels of transcripts in Chronic samples compared to Acute Baseline samples (Fig 2D and 2E). 13.3% and 13.4% of Signaling and Cellular Processes genes were significantly more highly expressed in Chronic Baseline or Enriched nutrient samples compared to Acute Baseline samples, whereas only 6.1% and 7.8% of Genetic Information Processing genes and 9.8% and 6.6% of Metabolism genes were significantly more highly expressed in Chronic Baseline and Chronic Enriched nutrient samples (Fig 2B and 2C). Thus, of these differentially expressed genes, signaling and cellular processes genes were disproportionately differentially expressed, with higher expression in Chronic Baseline and Chronic Enriched nutrient conditions, compared to genetic information processing and metabolism associated genes.

Several putative parasitism-associated genes were significantly differentially expressed in comparisons between Acute Baseline vs Chronic Baseline or Chronic Enriched nutrient samples and had significantly lower relative expression in Acute Baseline samples (Fig 2C). These included a T4SS gene (*virB10*) (CE [49.7 +/− 10.1], CB [40.0 +/− 3.6], AB [19.0 +/− 6.2] transcripts/sample), genes involved in amino acid transport (leu/ile/val and proline/betaine) (CE [6.3 +/− 1.2], CB [10.0 +/− 1.7], AB [2.3 +/− 2.1] transcripts/sample), cationic antimicrobial peptide (CAMP) resistance (CE [8.0 +/− 3.0], CB [8.3 +/− 3.1], AB [1.7 +/− 1.5] transcripts/sample), and a flagellar regulator (*flrC)* (CE [60.0 +/− 12.3], CB [58.3 +/− 16], AB [29.0 +/− 7.0] transcripts/sample). A toxin/antitoxin transcriptional regulator had significantly lower expression in Chronic Baseline nutrient samples compared to both Acute Baseline and Chronic Elevated nutrient samples (CE [5.0 +/− 1.0], CB [1.0 +/− 1.0], AB [7.0 +/− 2.6] transcripts/sample).

### Putative virulence factors are highly expressed in Chronic Baseline and Chronic Enriched nutrient treatments

Given that differentially expressed Signaling and Cellular Processes genes appear to disproportionately drive differences between ‘*Ca.’* A. rohweri transcriptomes, we wanted to more closely examine the specific function of genes within these categories. The alphaproteobacteria ‘*Ca.’* A. rohweri encodes a variety of membrane transport genes including ABC transporters, Major facilitator transporters (MFSs), Drug/Metabolite Transporters (DMTs) and a type 4 secretion system (T4SS). As a whole, membrane transport genes had significantly higher expression in Chronic Baseline and Chronic Enriched nutrient samples (average of 1,867 and 1,894 transcripts/sample) compared to Acute Baseline samples (average of 1,348 transcripts/sample) (two-way ANOVA with Tukey’s multiple comparison test, *P*<0.02) (Fig S3).

Hierarchical clustering analysis revealed that Acute Baseline samples had lower transporter expression compared to Chronic Baseline and Chronic Enriched nutrient samples on average (Fig S3). This observation was most striking for *eamA* genes, which encode a S-adenosylmethionine (SAM) transporter. When we looked only at expression of *eamA* genes, Acute Baseline samples clustered separately from Chronic samples (Fig 3A), suggesting these genes are particularly responsive to enriched nutrients.

**Figure 3.**
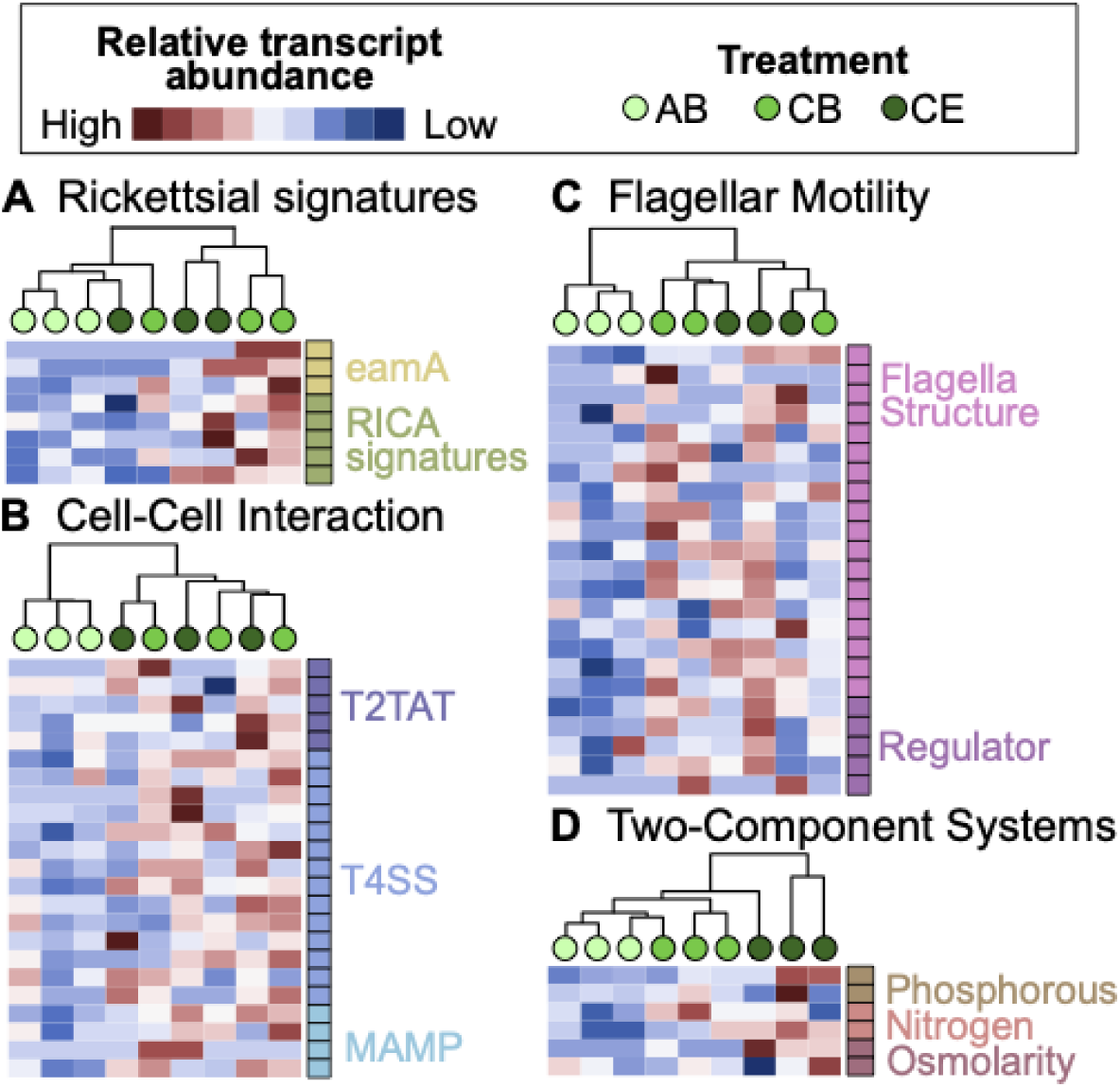
Rickettsial signatures, cell-cell interaction, motility, and two-component system genes are upregulated under nutrient enrichment. Hierarchical clustering analysis and heatmaps displaying relative transcript abundance for a subset of cell-cell interaction and environmental sensing genes. Each circle within the hierarchical clustering analysis represents a sample and circle color indicates the experimental treatment: Acute Baseline (AB, light green), Chronic Baseline (CB, medium green), Chronic Elevated nutrients (CE, dark green). Relative transcript abundance is scaled across samples for a given gene where genes with relatively highly transcript abundance are shown in red and those with relatively low abundance are shown in blue. Squares to the right of each heatmap indicate gene annotations. (A) Rickettsial signatures including 3 *eamA* genes, *proP, spoT, tlc1, mdlB,* and *gltP* (ordered top to bottom RICA signatures rows), (B) Cell-cell interaction genes: type II toxin anti-toxin system (T2TAT), type IV secretion system (T4SS), microbe/pathogen-associated molecular patterns (MAMP). (C) Motility associated genes. (D) Two-component system genes.

We next examined expression of ‘cell-cell interaction’ and ‘motility associated’ genes. Cell-cell interaction genes were significantly upregulated in Chronic samples (average of 1,474 and 1,276 transcripts/sample) compared to Acute Baseline samples (average of 804 transcripts/sample) (two-way ANOVA with Tukey’s multiple comparison test, *P*<0.004) (Fig 3B). Hierarchical clustering analysis revealed that for a subset of these genes, specifically those including a toxin/antitoxin system, a T4SS, prokaryotic defense strategies, and four microbe-associated molecular patterns (MAMPs), Acute Baseline samples had significantly lower expression (two-way ANOVA with Tukey’s multiple comparison test (*P<* 0.001)) and clustered separately from Chronic samples (Fig 3B, Table S2). Genes encoding flagellar motility and chemotaxis showed a similar trend (Table S2) and had significantly higher expression in Chronic samples (two-way ANOVA with Tukey’s multiple comparison test (*P*<0.0001)) (Fig 3C).

### Two component systems are expressed in a nutrient concentration and exposure duration-dependent manner and are phylogenetically congruent with one another

Two-component systems (TCSs) allow bacteria to sense changes in environmental stimuli and mediate an adaptive response, mainly through changes in gene expression, and TCSs frequently regulate virulence factors of pathogenic bacteria (Beier & Gross 2006). ‘*Ca.’* A. rohweri encodes three two-component systems to sense and respond to phosphorus (PhoR-PhoB), nitrogen (NtrY-NtrX), and osmolarity changes (EnvZ-OmpR). The three TCSs encoded by ‘*Ca.’* A. rohweri were significantly upregulated under chronic exposure to tank conditions and enriched nutrients in a dose-dependent manner (Fig 3D) (Fig 3D, two-way ANOVA with Tukey’s multiple comparison test (*P<*0.03)). This is interesting given that the nutrients elevated were N and P.

Given the limited experimental evidence determining the function of these two-component systems in Rickettsiales bacteria, we sought to assess the evolutionary relationships among them and Rickettsiales phylogeny. We constructed a series of maximum-likelihood phylogenetic trees using histidine kinase and response regulator amino acid sequences for each two-component system, as described in Speare *et al.,* (*30*). All three two-component systems were phylogenetically congruent to one another, suggesting a shared evolutionary history. To determine the evolutionary relationship of these two-component systems to Rickettsiales phylogeny, we constructed a consensus phylogenetic tree of all three two component-systems (six amino acid sequences total) and compared it to a 16S maximum likelihood tree (Fig S5). Congruence among distance matrices (CADM) analysis revealed that there was phylogenetic congruence between two-component systems and 16S (Fig 3E), suggesting these systems share a common evolutionary history.

## Discussion

Using experimental dosing, transmission electron microscopy, meta-trascriptomics, and phylogenetics, we showed here that the novel marine Rickettsiales bacterium, ‘*Ca.’* A. rohweri, exhibits unique parasitic transcriptional activity within disease-susceptible *A. cervicornis* tissue after chronic exposure to tank conditions containing enriched inorganic nutrients, indicating a potential shift in symbiotic status or life history during enrichment. Our evidence suggests that energy scavenging genes as well as those potentially involved in host habitat transition, specifically genes involved in signaling, putative virulence factors, motility, and nutrient import genes have elevated expression in chronic exposure to tank conditions containing enriched nutrients. Also ‘*Ca.’* A. rohweri two-component systems, which sense and respond to extracellular nitrogen, phosphorus, and changes in environmental osmolarity, are expressed in an experimental duration and nutrient dose-dependent manner and are phylogenetically congruent to one another and strain phylogeny. Moreover, the localization of these parasites during this experiment to epithelia mucocytes, along with these shifts in gene expression, demonstrate that these parasites are likely transitioning to a new life history stage and/or preparing for horizontal transmission.

### Working Model for parasitic *‘Ca.’* A. rohweri activity in A. cervicornis tissue

In combination with previous observations, our data support a model whereby chronic nutrient enrichment negatively impacts the coral host through the interactive effects of dinoflagellate and Rickettsiales activity (Fig 4). Endosymbiotic coral dinoflagellates respond positively to elevated nitrogen and phosphorus by increasing in population density (11, 27) and remaining mutualistic with their coral hosts (31–33). In response to environmental stress and/or dinoflagellate density (34–37), corals produce additional mucocytes to defend against sedimentation and invading microbes (38–40). Such mucocyte production opens additional ecological niches for ‘*Ca.’* A. rohweri to quickly infect and proliferate (Fig 4). Two-component systems expressed by ‘*Ca.’* A. rohweri sense these changes in the extracellular environment within *A. cervicornis* tissue. As ‘*Ca.’* A. rohweri cells infect new mucocytes, they experience an increase in osmotic stress resulting in increased *envZ-ompR* expression (Yuan et al 2011). In response to elevated nitrogen and phosphorus, as well as elevated ATP concentrations in uninfected relative to highly infected mucocytes, ‘*Ca.’* A. rohweri increases expression of host energy scavenging genes including *tlc1,* amino acid importers, and transporters, allowing ‘*Ca.’* A. rohweri to siphon and potentially deplete host resources. Increased expression of the *rvh* T4SS, toxin/antitoxin systems, MAMPs, and flagellar genes likely contribute to host attachment, infection, and protection from host defenses. The combined effects of these transcriptional shifts allow ‘*Ca.’* A. rohweri to dominate and destabilize disease susceptible *A. cervicornis* microbiomes (27), thereby significantly increasing coral disease susceptibility and mortality (18).

**Figure 4.**
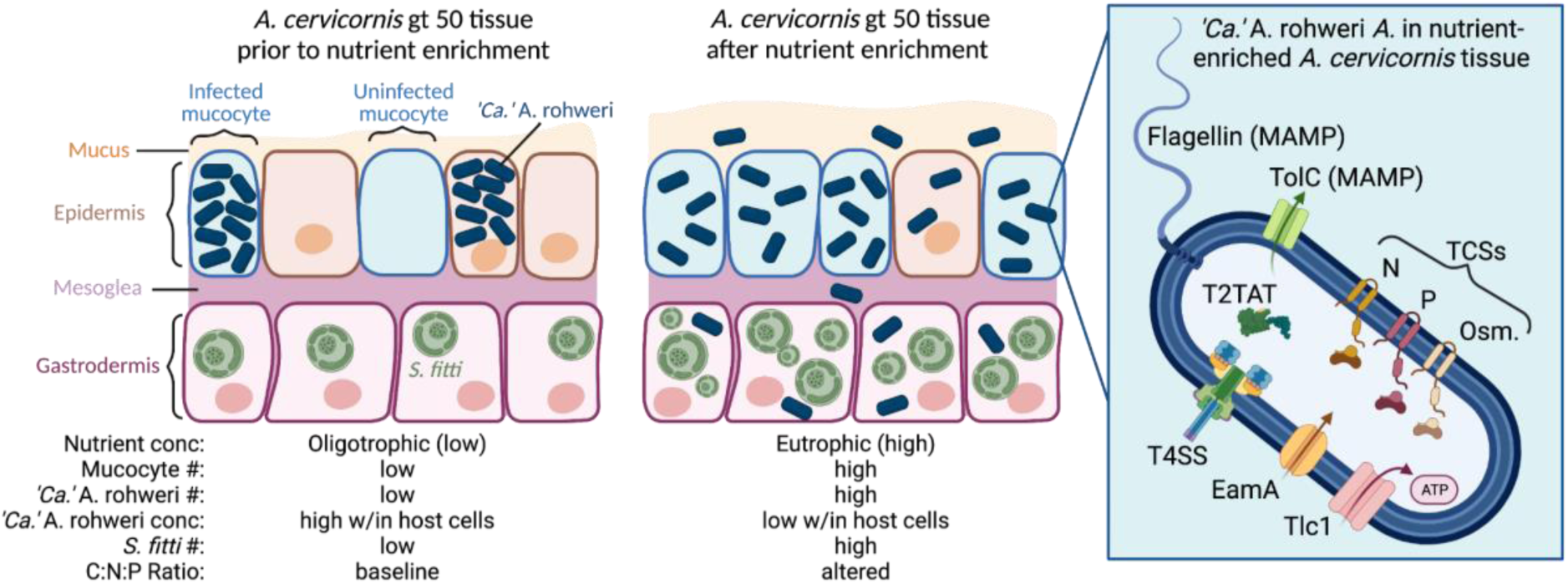
Conceptual model for the impact of nutrient enrichment on *‘Ca.’* A. rohweri activity and localization within *A. cervicornis* genotype ML-50 tissue. Diagrams on the left of the figure display predicted localization and concentration of ‘*Ca.’* A. rohweri (dark blue cells) and the endosymbiotic dinoflagellate *S. fitti* (green cells) within coral tissue prior to (left) or after (middle) nutrient enrichment experiment. Nutrient enrichment promotes mucocyte production (more light blue mucocytes), *S. fitti* density (green cells), and ‘*Ca.’ A. rohweri* abundance. Tlc1 expression data suggest that ‘*Ca.’* A. rohweri are present in lower concentrations within a given host cell. The schematic on the right displays gene products from differentially expressed genes with significantly higher expression in nutrient enriched conditions. Cells and structures are not displayed to scale; figure created using bioRender.com.

### Energy Acquisition & T4SS

Gene expression profiles of the PAMP *tlc1* support the predicted role of ‘*Ca.’* A. rohweri as a nutrient responsive coral parasite and provide insights into the energetic state of parasite and host cells under nutrient enrichment. Tlc1 functions as an ATP/ADP antiporter allowing Rickettsiales bacteria to siphon energy from host cells (41, 42). Mouse model studies examining *tlc* expression in lightly-vs heavily-Rickettsiales-infected host cells described lower levels of *tlc* mRNA in heavily-infected cells as compared to lightly-infected cells (43). Because Tlc1 can exchange ATP for ADP in both directions, lowering *tlc* expression would minimize Rickettsiales ATP efflux from the parasite back to the host when the energy pool is low in the host cytoplasm (44). ‘*Ca.’* A. rohweri infection correlates with reduced coral growth (9) and increased coral tissue loss (5), suggesting ‘*Ca.’* A. rohweri Tlc1 functions primarily to scavenge rather than provide ATP for infected cells. Higher expression of *tlc1* in our Chronic Baseline and Chronic Enriched nutrient enriched samples therefore suggest that ‘*Ca.’* A. rohweri ATP levels remain lower than ATP levels within the host cells they inhabit. If corals are, in fact, producing additional mucocytes, these may serve as new sites of infection, allowing parasite densities to stay relatively low and therefore increasing and/or maintaining high *tlc1* expression. Indeed, histological analysis of both healthy and diseased coral tissue has correlated the number of “aggregates” containing Rickettsiales-like organisms with worsening coral tissue conditions (23).

Elevated expression of the *rvh* T4SS, which is thought to translocate effector proteins into host cells (45), as well as toxin/antitoxin genes in concert with elevated *tlc1* expression, is also consistent with a rapidly expanding ‘*Ca.’* A. rohweri infection. Energy gained through additional Tlc1 activity may be used by the energetically costly T4SS apparatus to translocate molecules out of ‘*Ca.’* A. rohweri cells. Given the apparent intracellular nature of ‘*Ca.’* A. rohweri (22) Fig 1), which would allow direct translocation of molecules into coral mucocytes independently of the T4SS, it is currently unclear whether T4SS functions to translocate molecules across exclusively coral membranes or also into endosymbiotic dinoflagellate cells. Notably, expression of *virB10*, a component of the *rvh* T4SS that senses bacterial intracellular ATP levels to coordinate protein translocation (46), showed a particularly strong response to duration and concentration of nutrient enrichment. *tlc1* expression was significantly positively correlated with *virB10* expression, explaining 49.7% of the variation in *virB10* expression (Fig S4), suggesting that *virB10* expression is directly linked to Tlc1 activity. However, further manipulative experimentation is necessary to prove this relationship.

### Nutrient Sensing

The three two-component systems encoded by ‘*Ca.’* A. rohweri showed a dose-dependent response to the duration of exposure to and concentration of nitrogen and phosphorus, suggesting that like the majority of two-component systems described (47), these systems are controlled via positive feedback in response to nutrient enrichment. Given that ‘*Ca.’* A. rohweri lacks complete nitrogen metabolism pathways (22), elevated nitrogen sensed by the NtrY-X system may serve as a signal for elevated amino acid and/or sugar production by endosymbiotic dinoflagellates. Nitrogen enrichment is known to promote dinoflagellate symbiont proliferation (48–51) while phosphate is thought to have a lesser impact given that its availability is controlled by the coral host via active transport (52, 53). Therefore, dedicating the NtrY-X system as a sensor of dinoflagellate symbiont activity would prepare ‘*Ca.’* A. rohweri to quickly siphon photosynthates from the host or other members of the microbiome. Paired experimental data from our previous study, however, indicated that inorganic phosphorus rather than nitrogen, was the primary nutrient driving shifts in *Aquarickettsia* abundance (27). Thus, the high conservation of these and the EnvZ-OmpR two-component systems to one another and strain phylogeny supports the distinct, yet important function of each of these systems for ‘*Ca.’* A. rohweri fitness within host tissue.

‘*Ca.’* A. rohweri transcriptomes showed an unexpected increase in transporter gene expression under nutrient enrichment. Generally, nutrient depletion, rather than nutrient enrichment, triggers upregulation of nutrient acquisition systems, such as importers and/or biosynthesis pathways, allowing bacteria to effectively scavenge nutrients that may be scarce. However, a dissolved organic carbon (DOC) enrichment experiment revealed that coastal bacterioplankton upregulate gene expression of amino acid and sugar transporters in response to elevated DOC (54). Given that we currently know little about the environmental conditions that regulate Rickettsiales transporter expression, our data suggest that ‘*Ca.’* A. rohweri may upregulate transporter expression in response to chronic enriched nutrients. Alternatively, *‘Ca.’* A. rohweri may sense artificially lower levels of nitrogen and phosphorus than we measured in the aquaria. It is possible that dinoflagellate photosynthesis during enrichment alters the C:N:P ratio (55), i.e. the dinoflagellate symbionts become “greedy” and share less photosynthates with the coral host, such that ‘*Ca.’* A. rohweri sense artificially lower levels of nitrogen and phosphorus compared to carbon, resulting in upregulation of transporters. It is also possible that newly created mucocytes contain relatively low concentrations of nutrients, promoting *‘Ca.’* A. rohweri cells infecting newly created mucocytes to increase nutrient transport expression. Mucus secreted from coral mucocytes is enriched with high concentrations of endosymbiotic dinoflagellate photosynthates and derivatives of coral heterotrophic feeding (40, 56, 57), however whether this process occurs during or after mucocyte development, or after mucus expulsion from mucocytes requires investigation. This observation could be influenced by elevated nutrients at our Ambient Baseline treatment compared to offshore conditions.

### Potential Mechanisms of Horizontal Transmission

Given that ‘*Ca.’* A. rohweri is transmitted horizontally and that the water column is substantially less nutrient dense than coral tissue/mucus, elevated nutrient acquisition expression and movement into dense mucocytes could also indicate that ‘*Ca.’* A. rohweri is preparing to evacuate host tissues. For example, if ‘*Ca.’* A. rohweri were evacuated from host tissue into the water column, they would likely experience a significant decrease in extracellular nutrients that would theoretically favor upregulation of nutrient acquisition systems. Corals secrete large amounts of mucus from mucocytes under environmental stress (58) and ‘*Ca.’* A. rohweri are likely transmitted horizontally through mucus expulsion (25), as Rickettsiales-like organisms are commonly observed in *Acropora* mucocytes in both healthy and diseased hosts (59–61). Elevated expression of flagellar motility genes and T4SS and toxin/antitoxin genes may be necessary for ‘*Ca.’* A. rohweri to successfully transition to and infect new hosts. The *fliF* gene in particular, which is involved in membrane (M)-supramembrane (S) ring assembly, one of the first steps of flagellar biogenesis (62), had notably high expression under chronic nutrient enrichment. ‘*Ca.’* A. rohweri flagellar genes could confer a variety of functions enhancing virulence including motility, protein export, or adhesion (63, 64), however, a specific role for Rickettsiales flagella has not yet been described.

## Conclusion

In this study we provide evidence that *‘Ca.’* A. rohweri acts as a highly active, nutrient-responsive parasite within host tissue, and we propose a working model for the negative, synergistic effect of coral symbiont activity in response to nutrient enrichment. Our findings suggest *‘Ca.’* A. rohweri expresses key environmental sensing and virulence genes in *A. cervicornis* genotype ML-50 tissue and supports previous observations that *‘Ca’.* A. rohweri is transmitted horizontally between hosts, possibly via mucocytes and/or an unknown mechanism. Although we did not detect enough *‘Ca.’* A. rohweri transcripts in disease-resistant *A. cervicornis* genotype ML-7 transcriptomes for analysis, this reduced level of detectable transcription suggests that regardless of how this parasite is acting, it likely has a limited effect on the coral host in disease-resistant compared to disease-susceptible genotypes.

Further investigation should explore whether ‘*Ca.’* A. rohweri exhibits similar parasitic activity toward disease resistant *A. cervicornis* genotypes, where ‘*Ca.’* A. rohweri is only a minor constituent of the microbiome, 2.5% as compared to 89.7% of microbiomes in apparently healthy disease susceptible *A. cervicornis* (18). It is possible that ‘*Ca.’* A. rohweri elicits a similar response to enriched nutrients in disease resistant genotypes, however the host does not succumb to parasitic effects due to the low total abundance of ‘*Ca.’* A. rohweri within the microbiome. Alternatively, disease resistant host genotypes may prevent parasitic infection or parasite activity by modulating host-microbe interactions and intracellular conditions (65), effectively dampening the nutrient induced responses of the parasite. Although this work did not directly examine whether ‘*Ca.’* A. rohweri parasitizes both coral and dinoflagellate cells, symbiotic dinoflagellate abundance significantly increased in response to enriched nutrients in our samples (27), suggesting that the effect of parasitic activity toward *S. fitti,* if any, was minimal. However, whether ‘*Ca.’* A. rohweri negatively affects the dinoflagellate symbiont as well as the coral host warrants further study.

## Methods

### Experimental Design & Sample Collection

To test the impacts of nutrient enrichment on *A. cervicornis* health and microbiome community function, a six-week tank experiment was conducted as previously described (24, 27) at the Mote Marine Laboratory International Center for Coral Reef Research & Restoration (24°39’41.9"N, 81°27’15.5"W) in Summerland Key, Florida from April to June 2019. The experiment was conducted in 4.7 L flow-through, temperature-controlled aquaria with natural locally-sourced sea water from the Atlantic side of the Keys. Sand- and particle-filtered water was fed from header tanks to aquaria by powerheads fitted with tubing splitters (two tanks per powerhead) at a flow rate of 256.66 ± 43.89 mL per minute. Aquaria were located outdoors under natural light regimes with the addition of 75% shade cloth to account for shallow aquarium depth. Aquaria were divided between two flow-through seawater raceways (20 aquaria per raceway), which allowed for temperature regulation of individual aquaria. Raceway water was prevented from entering aquaria by maintaining water levels below in and outflow holes using a standpipe. Water temperatures were maintained at an average of 27.19 ± 0.6 °C. Temperature was controlled by a boiler and chiller using a dual heat exchanger system connected to header tanks and individual raceways. Header tank pH was stabilized at ∼8.0 by aeration and mixed via a venturi pump system. Nutrient levels in aquaria were elevated compared to conditions at the coral collection site (Mote Marine Laboratory’s in situ coral nursery in Looe Key) as intake pipes were located in coastal water instead of offshore reef water (Fig. S1). While ammonium and phosphate levels were similar to reef conditions, nitrate concentration was 4-fold higher in aquaria compared to Looe Key. Aquaria were cleaned every third day to prevent overgrowth of coral fragments with diatoms or algae.

Fragments (∼5cm) of *Acropora cervicornis* genotype ML-50 (Coral Sample Registry accession: fa13971c-ea34-459e-2f13-7bfddbafd327) were collected from the Mote Marine Laboratory *in situ* coral nursery in Looe Key in April 2019. This genotype was previously delineated via microsatellite genotyping and was found to have a high level of disease susceptibility (18, 19). Six fragments were housed in each aquarium. Prior to experimental manipulation, fragments were allowed to acclimate to aquarium conditions for seven days. Nutrient enrichment was performed four times a day for 42 days (six weeks). Flow in all aquaria was stopped for an hour following nutrient amendment. This resulted in an hour-long nutrient ‘pulse’ four times a day, followed by an hour of dilution and four hours of exposure to ambient conditions. Coral fragments were sacrificed for sampling at three time points throughout the experiment: prior to nutrient exposure (T0), after three weeks, and after six weeks; only samples from T0 and six weeks were used for transcriptome analysis. Using sterile bone cutters, tissue was scraped from each fragment (avoiding the apical tip) and added directly to 2 mL tubes containing 0.5ml DNA/RNA shield (Zymo Research) and Lysing Matrix A (MP Biomedicals, 0.5 g garnet matrix and one 1/4" ceramic sphere). Tubes were immediately preserved at −80°C until further processing. Total RNA was extracted from 500 μl of tissue slurry using the E.Z.N.A.® DNA/RNA Isolation Kit (Omega Bio-Tek) and then stored at −80°C until further processing.

### Scanning Electron Microscopy

Samples were processed for Scanning Electron Microscopy at Oregon State University. Samples were decalcified for five weeks with a 10% EDTA (pH 7) solution; the solution was replaced three to four times each week due to the formation of white deposits on coral tissue, presumably from the dissolved skeleton. After the skeleton was fully dissolved, the remaining tissue was fixed with Karvosky fixative (2% paraformaldehyde, 2.5% glutaraldehyde, 0.1 M buffer) overnight. Samples were then embedded in agar for post-fixation staining performed by Teresa Sawyer at the Oregon State University Electron Microscope Facility. Briefly, coral tissue was first rinsed with 0.1M sodium cacodylate buffer. Post fixation was conducted in 1.5% potassium ferrocyanide and 2% osmium tetroxide in deionized water. Samples then underwent T-O-T-O staining, uranuylacetate, and lead aspartate fixation. Samples were sequentially dehydrated in a range of increasing concentration acetone mixtures for 10-15 minutes: 10%, 30%, 50%, 70%, 90%, 100%, 100%. Finally, samples were infiltrated with Araldite resin and ultrathin sectioned. Images were collected on a FEI Helios Nanolab 650 in STEM mode at the Oregon State University Electron Microscopy Facility.

### Sequencing & Bioinformatics Analysis

Residual DNA contamination was removed from RNA isolates using the RQ1 RNase-Free DNase (Promega). Ribosomal RNA was removed using equal parts ‘plant leaf’, ‘human/mouse’ and ‘bacteria’ Ribo-Zero kits (Illumina). RNA quality and concentration were verified by BioAnalyzer (Agilent Technologies, Santa Clara, CA) and quantitative PCR, respectively. cDNA library prep and sequencing was performed at Oregon State University’s Center for Quantitative Life Sciences (CQLS) Core Laboratories with the HiSeq 3000 platform. Three biological replicates for each treatment were sequenced (n=9). Quality scores were calculated for each sequence using FastQC and MultiQC; low-quality scores (average score <20 across 5bp) were removed. Adapters were trimmed using bbduck (BBTools User Guide); successful trimming was confirmed using FastQC/MultiQC. Forward and reverse reads were then interleaved using reformat (BBTools User Guide), mapped to the ‘*Ca.’* A. rohweri genome using BowTie2, and counted using HTSeq-count. The limit of detection for each gene was one read per gene.

The vegan package in R was used to perform principal coordinates analysis (PCoA) using the Bray-Curtis dissimilarity index, PerMANOVA using the Adonis function, and beta-diversity using the permutest.betadisper function. Gene categorization was performed based on Kyoto Encyclopedia of Genes and Genomes Orthology (KO) designations. Differential expression analysis was performed through DeSeq2 and as described previously (66). Briefly, a contingency table was generated by comparing average transcript per sample abundances between treatments using a one-way analysis of variance (ANOVA) corrected for multiple comparisons using the false discovery rate (FDR). Volcano plots were generated by graphing the negative log 10 q-value and log 2-fold change between treatments. Only transcripts with a log 10 q-value of 0.05 and a log 2 fold change >|1| are considered statistically significantly differentially expressed (DE). Data were graphed in Graphpad Prism and edited for publication using Inkscape 1.0.

### Phylogenetic analyses

Multilocus two-component system (TCS) phylogenetic analyses were performed using the response regulator and histidine kinase for the three two-component systems encoded by ‘*Ca.’* A. rohweri: NtrY-NtrX, PhoR-PhoB, and EnvZ-OmpR. Published sequence data from the genomes of X Rickettsiales bacteria were collected into three separate TCS files and combined into a single concatenated sequence for each TCS (ordered histidine kinase, response regulator). Concatenated sequences were aligned (ClustalW) and phylogenetic reconstructions assuming a tree-like topology were created with MEGAX via maximum likelihood (ML). Gaps were treated as missing. The LG model with non-uniformity of evolutionary rates among sites may be modeled by using a discrete Gamma distribution (+G) with 5 rate categories and by assuming that a certain fraction of sites are evolutionary invariable (+I) was the most optimal evolutionary model. Tree inference was applied heuristically via the nearest-neighbor-interchange [NNI] method without a branch swap filter for 1,000 bootstrap replications. Phylogenetic trees were visualized with MEGAX and edited for publication with Inkscape 1.0.

## Supporting information

SI files for Speare et al In Prep 2023

## Acknowledgements and Funding Sources

We would like to thank the Florida Keys National Marine Sanctuary for authorizing the use of nursery-reared corals under permit FKNM-2015-163 and Erich Bartels at Mote Marine Laboratory for propagating and providing the corals for research. We would like to acknowledge the assistance of Dr. Abigail Clark, Dr. Emily Hall, Alexandra Fine, Chelsea Petrik, and Kyle Knoblock at Mote Marine Laboratory’s Elizabeth Moore International Center for Coral Reef Research and Restoration for expertise and logistical help. We thank Dr. Kalia Bistolas, Savanah Leidholt, and Dr. Hannah Epstein for helpful discussions. Work in the Vega Thurber and Muller labs was funded by an NSF Biological Oceanography grant (#1923836). L. Speare was supported as a Simons Foundation Awardee of the Life Sciences Research Foundation. J Grace Klinges was funded by an NSF Graduate Fellowship (#1840998-DGE).

