## Supplementary material for "Nutrient enrichment alters gene expression in ‘*Ca.’* Aquarickettsia rohweri, promoting parasite expansion and horizontal transmission": SI files for Speare et al In Prep 2023

**Supplemental Tables & Figures**.

**Table S1. Nutrient levels for the enrichment experiment.** Nutrient level was assessed by AutoAnalyzer at Mote Marine Laboratory in Sarasota Florida. Nitrogen was measured as dissolved nitrate-nitrite, and ammonia; phosphorus was measured at dissolved orthophosphate. Data are presented in micromolar concentrations.

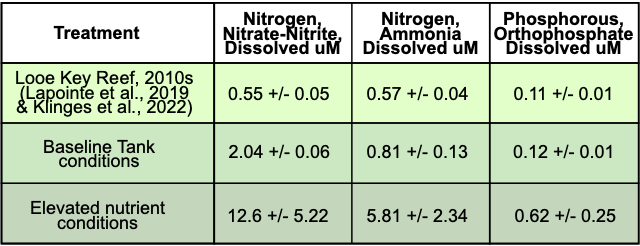

**Table S2. Transcripts per sample for cell-cell interaction and motility genes.** Table displays data shown in Figure 3B and 3C. Data are presented as the average number of transcripts per gene category across a given treatment (+/- standard deviation).

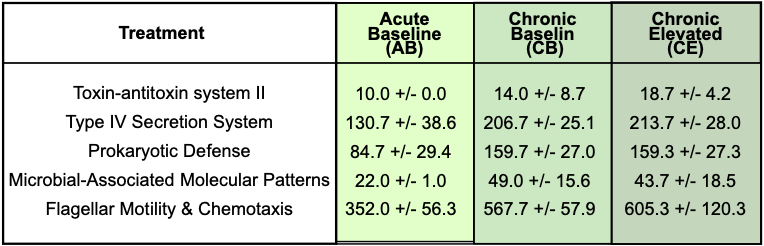

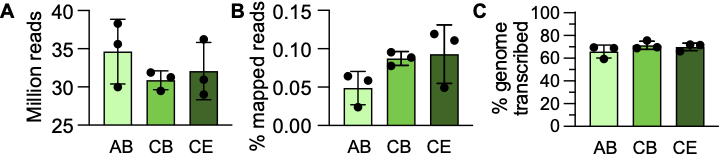

**Figure S1. Sequencing statistics for ‘*Ca.’* A. rohweri transcriptome.** Bar charts displaying (A) the number of reads for each sample after removing for low quality reads, (B) the percentage of reads that mapped to the ‘*Ca.’* A. rohweri transcriptome, and (C) the percentage of genes in the ‘*Ca.’* A. rohweri transcriptome that had at least one read. Bars are colored according to treatment: Acute Baseline (AB), Chronic Baseline (CB), or Chronic Enriched nutrients (CE). Values for each nutrient treatment were not statistically different from any other treatment in any of the panels shown above (One-way Analysis of Variance (ANOVA) with Tukey’s multiple comparison test, *P*>0.05).

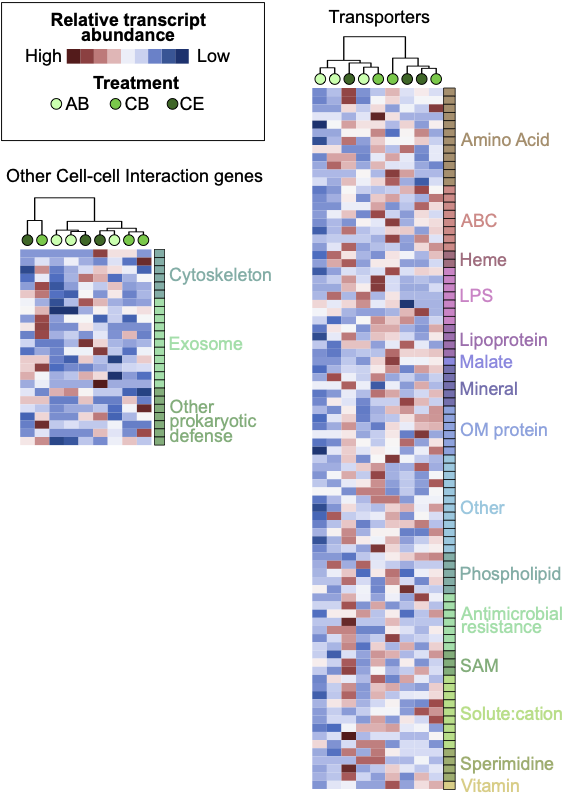

**Figure S2. Heatmaps and hierarchical clustering of other cell-cell interactions and transport genes.** Heatmaps display relative transcript abundance for a subset of cell-cell interaction and transport genes. Each circle within the hierarchical clustering analysis represents a sample and circle color indicates the experimental treatment: Acute Baseline (AB, light green), Chronic Baseline (CB, medium green), and Chronic Enriched nutrients (CE, dark green). Relative transcript abundance is scaled across samples for a given gene where genes with relatively highly transcript abundance are shown in red and those with relatively low abundance are shown in blue. Squares to the right of each heatmap indicate gene annotations.

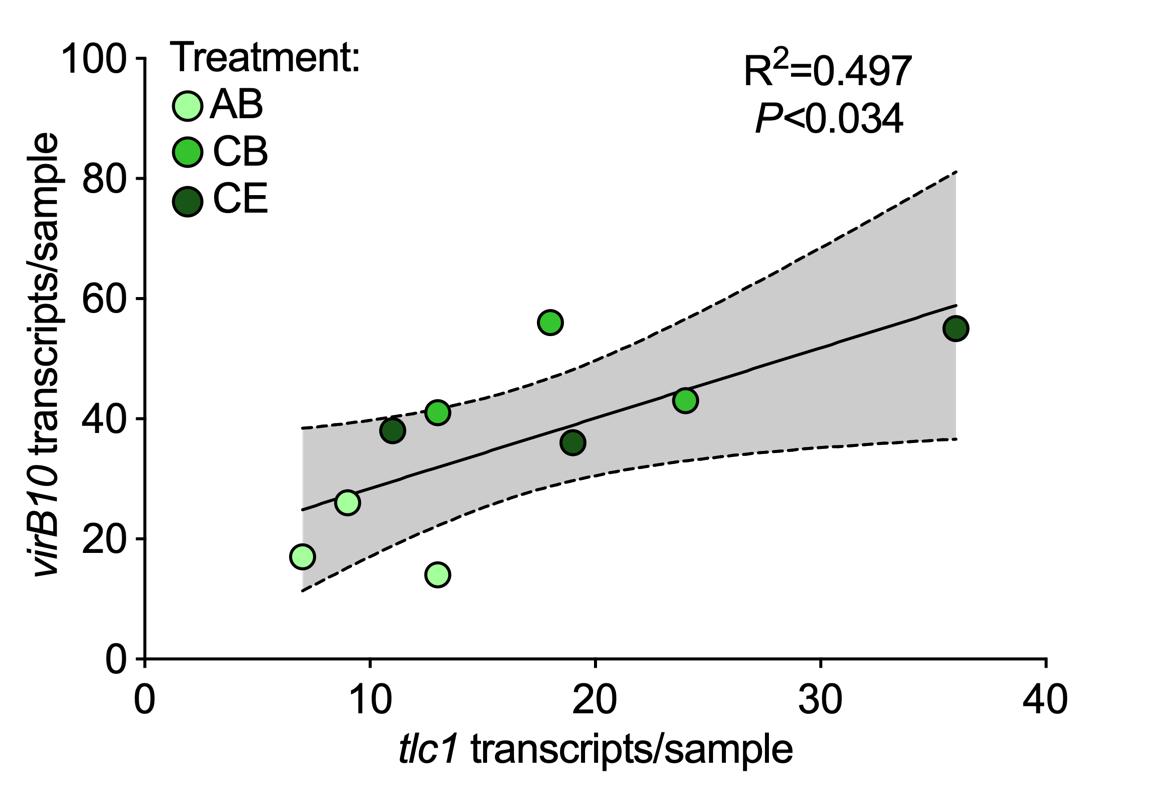

**Figure S3. Linear regression analysis reveals *virB10* expression is affected by *tlc1* expression in ‘Ca.’ *A. rohweri.*** Linear regression analysis between *tlc1* and *virB10* shows that *tlc1* expression explains 49.7% of the variation in *virB10* expression (simple linear regression, *P<*0.034)*.* Symbol color indicates the sample treatment: Acute Baseline (AB), Chronic Baseline (CB), and Chronic Enriched nutrients (CE). Gray area indicates 95% confidence interval.
